# Patient-Derived *PSEN1* Cerebral Organoids Revealed Parallel Development of Amyloid-β Accumulation and Network Dysfunction

**DOI:** 10.64898/2026.04.02.707990

**Authors:** Andrijana Angelovski, Hana Hribkova, Jiri Sedmik, Barbora Liscakova, Olga Svecova, Sona Cesnarikova, Katerina Amruz Cerna, Veronika Pospisilova, Martin Kral, Martina Kolajova, Petr Klimes, Dasa Bohaciakova, Marketa Bebarova

## Abstract

Alzheimer’s disease (AD) is a neurodegenerative disorder characterised by progressive dementia, brain atrophy, and ultimately death. Using cerebral organoids derived from human induced pluripotent stem cells (hiPSCs) carrying the familial *PSEN1* A246E variant, we investigated the temporal relationship between amyloid-β (Aβ) dysregulation and spontaneous neuronal activity. Multielectrode array recordings from the differentiation day 60 (DD60) to at least DD130 revealed that AD organoids exhibited transient hyperexcitability and hypersynchrony compared with wild-type (WT) controls, followed by a gradual decline in activity. During the enhanced excitability stage, both elevated Aβ42/40 and Aβ aggregate size showed positive correlations with the percentage of active electrodes and the global synchrony index (GSI) in AD organoids. These findings indicate that Aβ dysregulation might contribute to transient network hyperexcitability in early AD. The results also suggest that patient-derived cerebral organoids may serve as a translational model to examine early network dysfunction and inform future investigations of potential Aβ-induced changes in excitability during the preclinical stages of AD.

## 1. Introduction

Alzheimer’s disease (AD) is a progressive neurodegenerative disorder characterised, among other features, by the accumulation of amyloid-β (Aβ) plaques (DeTure & Dickson, 2019). Variants in the presenilin-1 (*PSEN1*) gene, which encodes the catalytic subunit of γ-secretase, are the most common cause of familial AD (fAD; Bagaria *et al*., 2022). These variants alter amyloid precursor protein (APP) processing, shifting Aβ production in favour of longer, less soluble Aβ peptides, including Aβ42, which leads to an increased ratio of Aβ42 to Aβ40 (Aβ42/Aβ40) and the formation of Aβ deposits in the brain. Nowadays, more than 300 *PSEN1* variants have been identified, yet they are associated with a broad spectrum of clinical phenotypes (Bagaria *et al*., 2022). Aβ42 may cause neuronal hyperexcitability, leading to overt seizures, impaired network oscillations, and epileptiform activity (Sciaccaluga *et al*., 2021). Several underlying mechanisms have been reported, including Aβ-dependent impairment of glutamate reuptake, dendritic structural degeneration, dysfunction of inhibitory interneurons, and aberrant modulation of ion channels (Busche & Hyman, 2020; Targa Dias Anastacio *et al*., 2022). In this context, *PSEN1* variants are particularly important as they provide insight into the early molecular mechanisms of altered excitability.

Although animal models have improved our understanding of AD pathogenesis, translating these findings into human therapy is significantly limited by species-specific differences. As a result, the application of human cerebral organoid models for AD research has been documented as a critical tool in several recent studies (*e.g.,* Shimada *et al*., 2022; Ji *et al*., 2025). These 3D neuronal/stem cell culture systems capture the complexity of the human brain *in vivo* more accurately, enabling the study of the functional roles of genetic variants, screening for potential therapeutics, and investigating mechanisms of disease initiation and progression *in vitro* (Barak *et al*., 2022; Vanova *et al*., 2023). While patch-clamp techniques can be used to study the electrophysiology of single neurons within cerebral organoids (Landry *et al*., 2023), multielectrode array (MEA) systems are better suited for investigating network-level activity (Angelovski & Bébarová, 2025). MEA recordings enable prolonged, repeated monitoring of electrical activity in organoids, allowing for the assessment of the relationship between electrophysiological properties and cellular or morphological features (Chung *et al*., 2022). Several studies using organoid-MEA platforms have already reported abnormal or increased network activity (Chen *et al*., 2021; Ghatak *et al*., 2021).

As previously mentioned, various mechanisms have been proposed through which Aβ oligomers can induce hyperexcitability in AD. However, the temporal relationship between Aβ dynamics and neuronal electrical activity remains poorly understood. To address this, we investigated how changes in Aβ levels correlate with spontaneous neuronal activity over time using previously characterised cerebral organoids derived from a fAD patient carrying a *PSEN1* A246E variant (abbreviated as AD) and an unrelated healthy donor, wild type (WT; Vanova *et al*., 2023). Our study reveals a positive association between Aβ42/Aβ40, Aβ aggregate size, and spontaneous electrical activity during the early stages of AD.

## 2. Methods

### 2.1 Human Induced Pluripotent Stem Cell Culture and Cerebral Organoid Differentiation

Human induced pluripotent stem cells (hiPSCs) were previously generated from fibroblasts (Raska *et al*., 2021). Two hiPSC lines were used in this study, one from a healthy donor (MUNIi009-A; RRID:CVCL_A4PH; abbreviated as WT) and one from a patient carrying the *PSEN1* A246E variant (MUNIi006-A; RRID:CVCL_A4PE; abbreviated as AD). hiPSCs were cultured on Matrigel-coated plates (hESC-qualified; Corning) in mTeSR1 medium (STEMCELL Technologies) supplemented with 1% ZellShield (Minerva Biolabs) under standard conditions (37 °C, 5% CO₂). Colonies were passaged every 5–7 days using manual passaging and routinely tested for mycoplasma.

Cerebral organoids were generated using a modified version of the protocol by Lancaster *et al*. (2013) and Lancaster and Knoblich (2014) as described previously (Vanova *et al*., 2023). Briefly, hiPSCs were collected using Accutase (Thermo Fisher) and aggregated into embryoid bodies on the differentiation day 0 (DD0; 3,500 cells per 96-wp well in 150 µl of mTeSR1 medium supplemented with 50 μM Y-27632 (Selleckchem)). On DD2, fresh mTeSR1 without Y-27632 was added, followed by daily feeding with Neural Induction Medium from DD3 till DD9. On DD9, organoids were embedded in 7 μl Geltrex droplets (A1413302; Thermo Fisher) and moved to Petri dishes with Cerebral Organoid Differentiation Medium without vitamin A. From DD21, organoids were maintained in Cerebral Organoid Differentiation Medium with vitamin A, and on DD27, they were transferred to an orbital shaker. On DD52-DD55, organoids were moved to MEA cultivation medium containing BrainPhys (STEMCELL Technologies) to support synaptic activity. Organoids were then maintained until DD130, with media changed every 2–3 days, and monitored for size, shape, and neuroepithelial organisation. All media compositions are summarised in **Supplementary Table S1**.

### 2.2 Multielectrode array recording and analysis

Spontaneous electrical activity of the cerebral organoids was recorded at 37 °C using the multielectrode array (MEA) MEA2100-Lite-System in combination with 60-electrode planar MEAs 60MEA-200/30iR-Ti and acquisition software Multi Channel Experimenter. The sampling rate was set to 25 kHz, with a high-pass filter of 10 Hz and a low-pass filter of 12.5 kHz (Multi Channel Systems GmbH, Reutlingen, Germany). The recordings were performed in MEA cultivation medium (**Supplementary Table S1**) for 15 minutes, twice a week, from DD60 to at least DD139, using a total of 15 AD and 16 WT cerebral organoids (3 independent differentiations, with at least 5 organoids per differentiation).

For analysis, a 10-minute interval between the 5^th^ and 15^th^ minute of each recording session was used. The raw signals were exported in the native Multichannel Systems format and converted into EDF format. After filtering the raw signal using a 300–3000 Hz band-pass filter to remove the noise, spikes representing multi- and single-unit activity were detected using the MEA-Tools Python Library (https://github.com/dbridges/mea-tools) and custom-made post-processing steps (the codes available here: https://gitlab.com/bbeer_group/research/mea/mea_sigproc.git). The threshold for spike detection was set at a minimum of ±6.5 standard deviations (SD) from the noise level. We discarded spikes that were considered artefacts due to their occurrence at the same time in more than 50% of electrodes. Using these detections, the following parameters were evaluated using Origin 2024 (OriginLab Corporation) and MATLAB R2023b (MathWorks):

i. spike parameters: active electrodes (%), spike frequency (Hz) (cut-off: ≥ 0.1 Hz), spike amplitude (µV) (cut-off: ≥ –6.5 SD of the baseline noise), interspike interval (ISI, s), and ISI coefficient of variation (ISIcv);
ii. burst parameters: active electrodes (%), burst count, burst frequency (bursts/min), interburst interval (IBI, s) (cut-off: ≥ 0.1 s), IBI coefficient of variation (IBIcv), burst duration (s) (cut-off: ≥ 0.1 s), intraburst spike number (cut-off: ≥ 5 spikes), and intraburst spike frequency (Hz);
iii. synchrony: the global synchrony index (GSI) calculated from detected spikes across active electrodes - A python script describing electrode synchronisation based on spike timing was written according to the information and methods proposed in literature (Patel *et al*., 2012; Eisenman *et al*., 2015). This includes necessary calculations, such as the phase transformation of spikes into phase time series and the quantification of pairwise synchronisation using a phase synchronisation matrix. GSI is then calculated as the strongest signal (given by the largest eigenvalue of the matrix) subtracted by 1 to remove the baseline self-correlation and scaled relatively to the number of active electrodes subtracted by 1 to normalise the index to the range [0,1]. If GSI = 0, then signals are independent, whereas if GSI = 1, then signals are fully synchronised.

### 2.3 Immunohistochemistry

Selected organoids were fixed in 4% paraformaldehyde and cut into 200 µm-thick sections using a Leica VT100S vibratome (Leica Biosystems, Germany). For immunohistochemistry, a homemade fructose–glycerol clearing agent was used according to Dekkers *et al*., 2019. In brief, sections were placed into a Greiner 96-well plate and blocked in 0.5% Triton X-100 and 5% bovine serum albumin (BSA) in phosphate-buffered saline (PBS). Primary antibodies were diluted in the blocking buffer at a ratio of 1:200 and incubated for 5 days on a shaker at 4 °C. The incubation was followed by a wash with 0.5% Triton X-100 in PBS. Secondary antibodies, diluted 1:400 in the blocking buffer, were applied overnight on the shaker at 4 °C. After washing, the fructose–glycerol clearing solution was used for microscopic observation and long-term storage. Sections were captured using a Zeiss LSM800 confocal microscope (Carl Zeiss, Germany). Final images were processed using Zen Blue (Carl Zeiss, Germany) and Imaris analysis software (Oxford Instruments, United Kingdom). List of primary and secondary antibodies is as follows: Aβ (Cell Signaling, 8243), SOX2 (Cell Signaling, 3579), PAX6 (Cell Signaling, 60433), NeuN (Millipore, MAB377), CTIP2 (Cell Signaling, 12120), MAP2 (Millipore, AB5543), TUJ (Aves Labs, TUJ-0020), Donkey Anti-Chicken AF647 (Jackson ImmunoResearch, 703-606-155), Donkey Anti-Mouse AF488 (Thermo Fisher Scientific, A-21202), Donkey Anti-Rabbit AF568 (Thermo Fisher Scientific, Cat# A-10042), and Donkey Anti-Rabbit AF647 (Thermo Fisher Scientific, A-31573).

Samples of sections of cerebral organoids were imaged with the inverted microscope Zeiss Axio Observer.Z1 with confocal unit LSM 800, equipped with solid state lasers (405, 488, 561, 640 nm), using Plan-Neofluar 5x/0.16 AIR objective, Plan-Neofluar 10x/0.30 AIR, and ZEN Blue software (Zeiss). Images with 0.618 x 0.618 x 20 µm (5x) and 0.329 x 0.329 x 4.2 µm (10x) pixel size were acquired using GaAsp PMT detectors. The acquisition parameters of detectors for Alexa Fluor 488, 568, and 647 were: 498-553 nm, 565-617 nm, and 656-700 nm (emission wavelength range) and 1.5 µs (pixel dwell time). Scan mode was set to frame, and the pinhole was set to 1AU. The line average of 2 was applied to all channels. To evaluate Aβ accumulation in thick sections of cerebral organoids, we used commercially available software Imaris version 9.8.2 (Bitplane). The detection of individual Nuclei and Aβ plaques was performed in Imaris using the “Surface” module. The estimated parameters included the volume of Aβ and nuclei. Parameters were automatically quantified using the Imaris software. Data were analysed and plotted using GraphPad Prism version 9 (GraphPad Software).

### 2.4 Enzyme-Linked Immunosorbent Assay (ELISA) for Aβ40 and Aβ42

Conditioned media from cerebral organoid cultures (1 organoid in 12-wp well with 0.8-1.2 ml of Essential 6 Medium (Thermo Fisher)) were collected at defined time points, centrifuged at 200 × *g* for 5 minutes to remove debris, and stored at –80 °C until analysis. Aβ40 and Aβ42 levels were measured with Amyloid beta 40 Human ELISA Kit (KHB3481, Thermo Fisher) and Amyloid beta 42 Human ELISA Kit, Ultrasensitive (KHB3544, Thermo Fisher) according to the manufacturer’s instructions. Samples and standards were run in duplicates, and absorbance was measured at 450 nm using a microplate reader. Concentrations were calculated from a standard curve and normalised to media volume and total protein content per organoid, as determined by DC Protein Assay (Bio-Rad) from parallel samples.

### 2.5 Quantitative Real-Time PCR (qPCR) Analysis of Neuronal Markers

Total RNA was isolated from 5-8 organoids with RNA Blue reagent (Top-Bio) according to the manufacturer’s instructions. The isolated RNA was reverse transcribed to cDNA using Transcriptor First Strand cDNA Synthesis Kit (Roche) according to the manufacturer’s instructions. qPCR was performed from the cDNA samples using LightCycler 480 SYBR Green I Master kit (Roche) on LightCycler 480 II (Roche). Gene-specific primers are listed in **Supplementary Table S1**. Ct values were calculated using the automated Second Derivative Maximum Method in LC480 software (Roche). Relative expression levels were calculated using the ΔΔCt method and normalised to the housekeeping gene *GAPDH*. Each sample was run in technical triplicate, and melting curve analysis was performed to confirm the specificity of amplification.

### 2.6 Data Analysis and Visualisation

GraphPad Prism version 9 (GraphPad Software) was used for statistical evaluation and graphical representation. Graphs and summary statistics were generated in Prism. Schematic illustrations and graphical summaries of experimental workflows were created using BioRender.com. Figures were assembled in Adobe Illustrator for final layout and labelling.

For immunohistochemistry analysis, Aβ aggregates were identified and quantified using Imaris software. To consider the statistical significance of differences in data showing the Aβ aggregate size and secreted Aβ40 and Aβ42, ANOVA with Šidák’s post-hoc test was performed; *P* < 0.05 was considered statistically significant.

Considering the rejection of the normal distribution in most MEA datasets (as indicated by the Shapiro-Wilk test), the data are presented as boxplots, including all measured points, and show the median and whiskers representing the minimal and maximal values. Differences between groups were evaluated using non-parametric tests, namely the Mann-Whitney test and Kruskal-Wallis test with Dunńs post hoc multiple comparison test as mentioned in the figure legends; *P* < 0.05 was considered statistically significant. To investigate possible correlations between electrophysiological and biochemical parameters, Spearman’s rank-order correlation was used (*r*_s_ - the Spearman correlation coefficient).

## 3. Results

### 3.1 Cerebral organoids differentiated from hiPSCs carrying the *PSEN1 A246E* variant accumulate AD-like pathology

To initiate our experiments, we generated cerebral organoids from hiPSCs carrying the early-onset familial AD-causing variant *PSEN1 A246E* (AD) and WT non-demented controls. Organoids were differentiated and matured over time to assess neurodevelopment, Aβ accumulation, and associated pathophysiological features. Immunofluorescence staining confirmed successful organoid formation and neuronal differentiation in both genotypes, demonstrating organised neuroepithelial structures and expression of progenitor markers (SOX2, PAX6), as well as cortical neuronal markers MAP2, NEUN, TUJ1, and CTIP (**Figure 1A**), consistent with a cortical-like identity.

**Figure 1:**
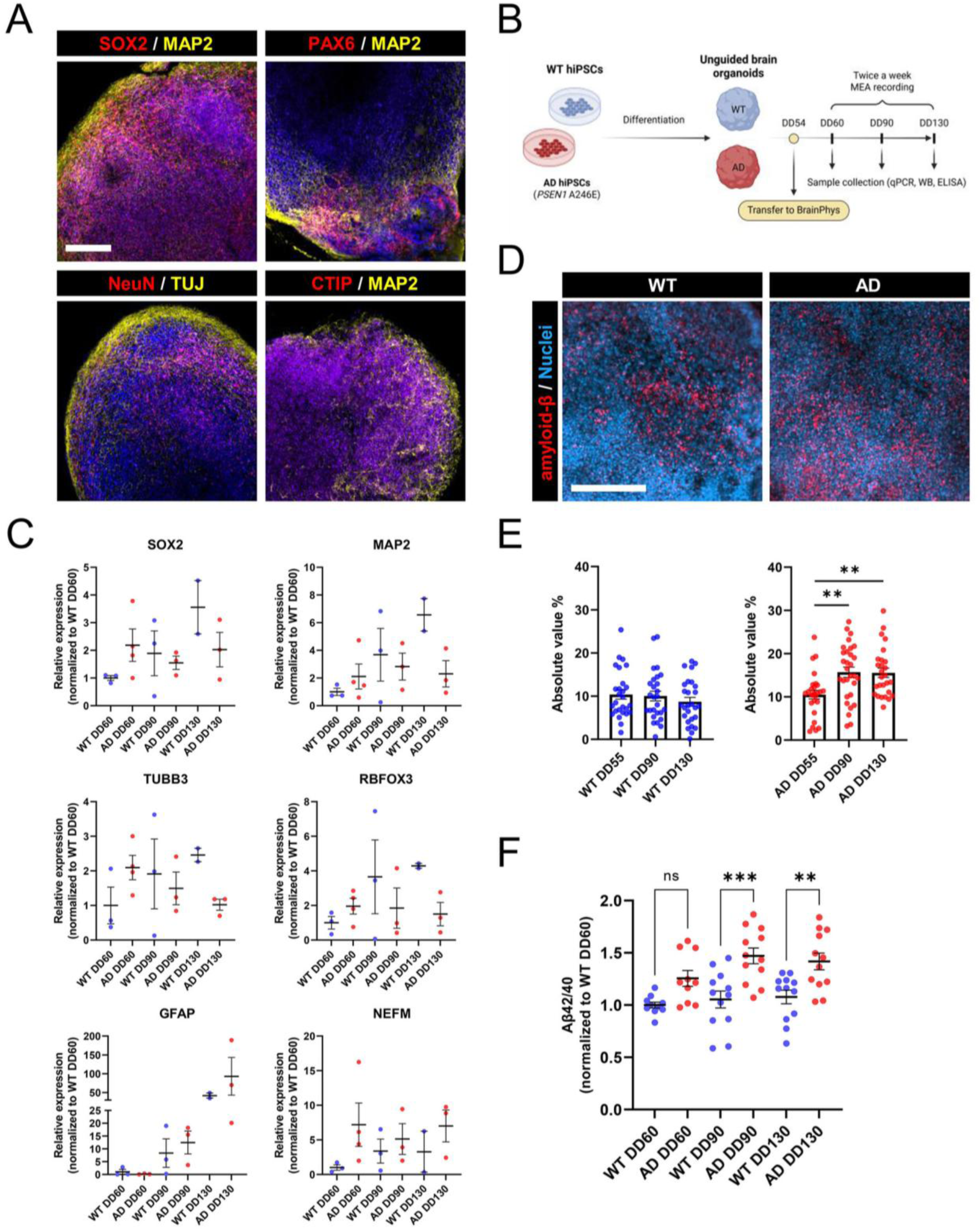
Experimental model and characterisation of Alzheimer’s disease-related phenotypes in cerebral organoids derived from hiPSCs carrying the *PSEN1* A246E variant. **A:** Representative immunofluorescence image of a wild-type (WT) cerebral organoid section showing well-organised neuroepithelial zones and expression of key neuronal markers, including SOX2, PAX6, NeuN, CTIP2, MAP2, and TUJ, confirming neural differentiation and cortical identity at differentiation day 90 (DD90). Scale bar = 200 µm. **B:** Experimental design: cerebral organoids were generated from WT and AD iPSC lines and harvested at differentiation days 60, 90, and 130 (DD60, DD90, and DD130) for molecular, histological, and biochemical analyses. **C:** qPCR analysis of selected neural stem cell, neuronal, and glial markers in DD60-DD130 organoids. Results are normalised to mRNA expression in WT DD60. Each dot represents a pooled sample of 5–8 organoids (n = 2-4 independent experiments) and the lines represent mean ± SEM. **D:** Representative immunofluorescence image showing amyloid-β (Aβ) accumulation in AD organoid section at day 90 (DD90) using the D54D2 antibody. Scale bar = 200 µm. **E:** Quantification of intracellular Aβ signal intensity in WT and AD organoids across all timepoints, revealing progressive accumulation in the AD model. ANOVA with Šidák’s post-hoc test was performed. \*\**P* < 0.01 **F:** ELISA of Aβ40 and Aβ42 secreted into organoid cultivation media. Aβ42/Aβ40 normalised to WT DD60 is shown. Each dot represents an individual organoid (n = 10–12; 3 independent experiments) and the lines represent mean ± SEM. ANOVA with Šidák’s post-hoc test was performed. \**P* < 0.05; \*\**P* < 0.01; \*\*\**P* < 0.001

Subsequently, to systematically assess disease progression, organoids were collected at three time points: DD60, DD90, and DD130 (**Figure 1B**). These time points were chosen to capture various stages of neuronal maturation and potential Aβ pathology. Organoids were then subjected to molecular, histological, and biochemical analyses at each stage to evaluate genotype-dependent effects. We first examined neuronal development using quantitative PCR for several neurodevelopmental and neuronal markers. Both WT and AD organoids expressed similar, relatively stable levels of neural stem cell and neuronal markers (SOX2, MAP2, TUBB3, and RBFOX3 (NeuN)), while the levels of astrocyte marker GFAP increased in later stages of differentiation. Notably, AD organoids showed slightly increased NEFM expression compared to WT, suggesting potential early alterations in neuronal integrity in AD organoids (**Figure 1C**).

Importantly, to assess hallmark AD pathology, we performed immunofluorescent staining for Aβ in organoid sections. In AD organoids, we observed focal accumulation of Aβ signal at later stages of differentiation (DD90), while WT organoids showed minimal signal (representative picture is shown in **Figure 1D**). Quantification of the Aβ immunoreactivity across timepoints confirmed a progressive increase in AD organoids, in contrast to stable levels observed in WT controls (**Figure 1E**).

Finally, to complement these findings, we measured secreted Aβ species in organoid-conditioned media using ELISA. AD organoids exhibited a consistently elevated Aβ42/Aβ40 at all analysed timepoints (DD60, DD90, and DD130) compared to WT organoids, supporting the known pathogenic shift in Aβ processing driven by *PSEN1* variants (**Figure 1F**). Together, these data validate our cerebral organoid model as a robust system for capturing early AD-related phenotypes, including altered neuronal gene expression and progressive Aβ accumulation, driven by the *PSEN1 A246E* variant.

### 3.2 Different patterns of electrical activity in AD and WT organoids

To examine the development of spontaneous network activity, we recorded extracellular signals from WT and AD organoids between DD60 and DD139 in 3 independently differentiated batches (*n* = 16/3 WT and 15/3 AD organoids), as described in the Methods section. Representative MEA recordings in a single electrode and respective heatmaps of spike counts within 10-minute recordings in all electrodes in representative WT and AD organoids on DD60, DD75, DD90, DD105, and DD120 are illustrated in **Figure 2**. Data show that electrical activity gradually increased and then decreased in AD organoids during the recording period, but it remained stable or even decreased and was substantially lower in WT organoids. These opposing temporal profiles indicate divergent maturation patterns of neuronal network dynamics in AD *versus* WT organoids.

**Figure 2:**
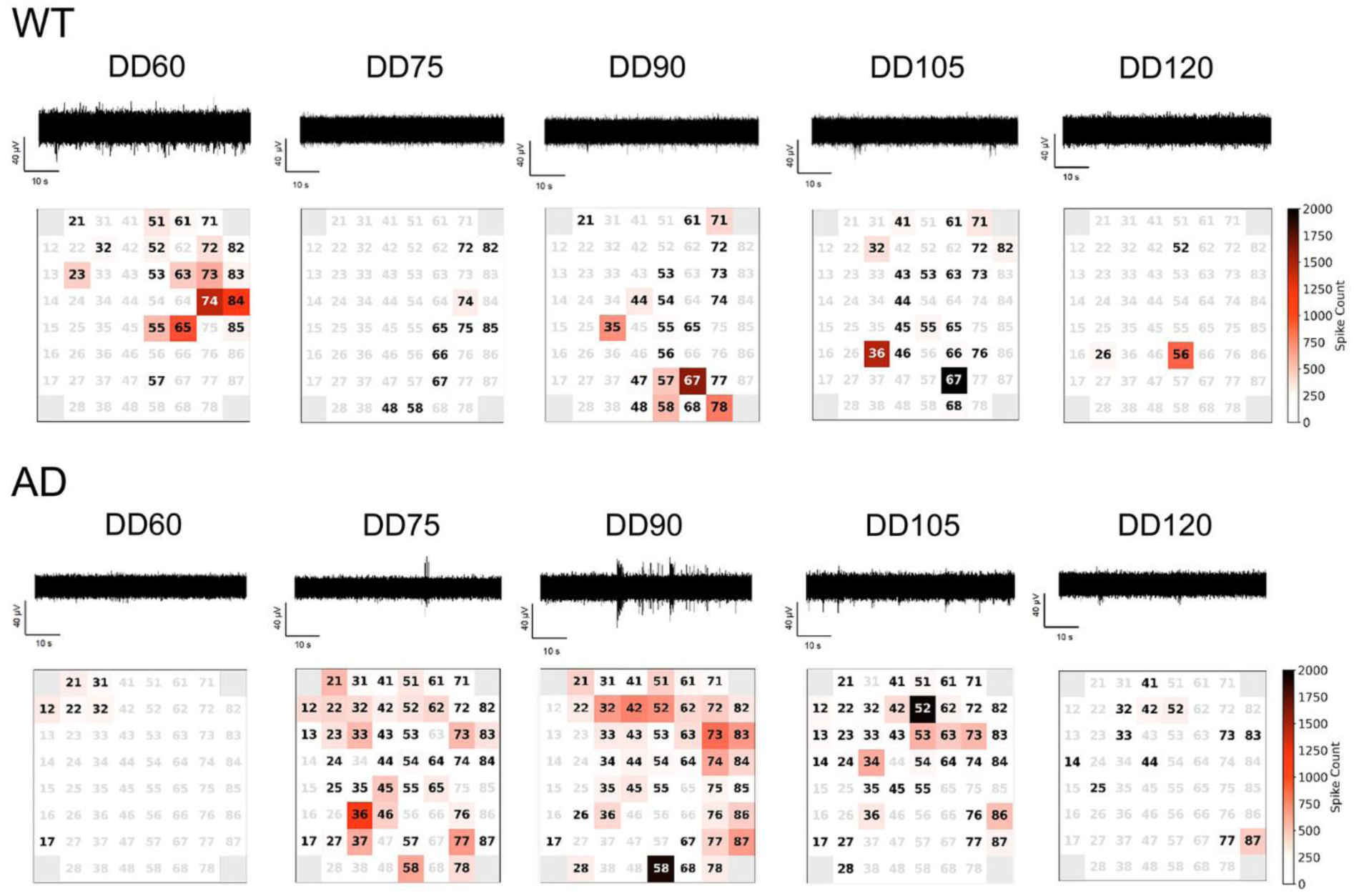
Electrical activity in WT and AD cerebral organoids recorded by the multielectrode array (MEA) technique. Representative MEA recordings at a single electrode (electrode numbers 55 and 42 in WT and AD, respectively) and respective heatmaps showing spike counts during 10-min MEA recordings at DD60, DD75, DD90, DD105, and DD120 in WT (up) and AD (down) organoids; grey electrode numbers – no electrical activity detected.

To better understand how neuronal firing properties differ between WT and AD organoids, we analysed spike parameters, reflecting the number and intensity of action potential–like events detected across electrodes (**Figure 3**). The percentage of active electrodes, an indicator of the number of neurons participating in network activity, remained relatively low and stable in WT organoids throughout the observation period. In contrast, AD organoids displayed a pronounced increase, peaking between DD72 and DD82 (**Figure 3A**). The percentage of active electrodes was significantly higher in AD organoids compared to WT at multiple time points, including DD72, DD75, DD79, DD82, DD98, DD100, and DD105 (\**P* < 0.05, \*\**P* < 0.01, and \*\*\**P* < 0.001). Additionally, the variability was also considerably higher in AD (**Figure 3A**). At the peak of the activity in AD organoids, the difference in the percentage of active electrodes was highly significant between AD and WT (\*\*\**P* < 0.001; **Figure 3B**). The spike frequency and interspike interval coefficient of variation (ISIcv) were comparable between AD and WT organoids across most time points, including the peak activity period (**Figures 3C, 3D, 3E, and 3F**). The spike amplitude showed significantly higher values in AD organoids compared to WT at DD60 and DD67 and during the peak of activity (\**P* < 0.05 for both; **Figures 3G** and **3H**). To consider differences in connectivity and, thus, in synchronisation of WT and AD organoids, the global synchrony index (GSI) was evaluated (for details, see Methods). AD organoids exhibited significantly higher GSI values than WT, particularly in the first 40 days of the recording period (*i.e.*, from DD60 to DD100; \**P* < 0.05, \*\**P* < 0.01, and \*\*\**P* < 0.001; **Figure 3I**). During the three weeks of the highest electrical activity in AD organoids, the significance of the difference in GSI was high (\*\*\**P* < 0.001; **Figure 3J**). Together, these findings indicate that AD organoids recruit more neurons into network activity and display stronger synchrony than WT, while the firing rate of individual neurons remains largely unchanged.

**Figure 3:**
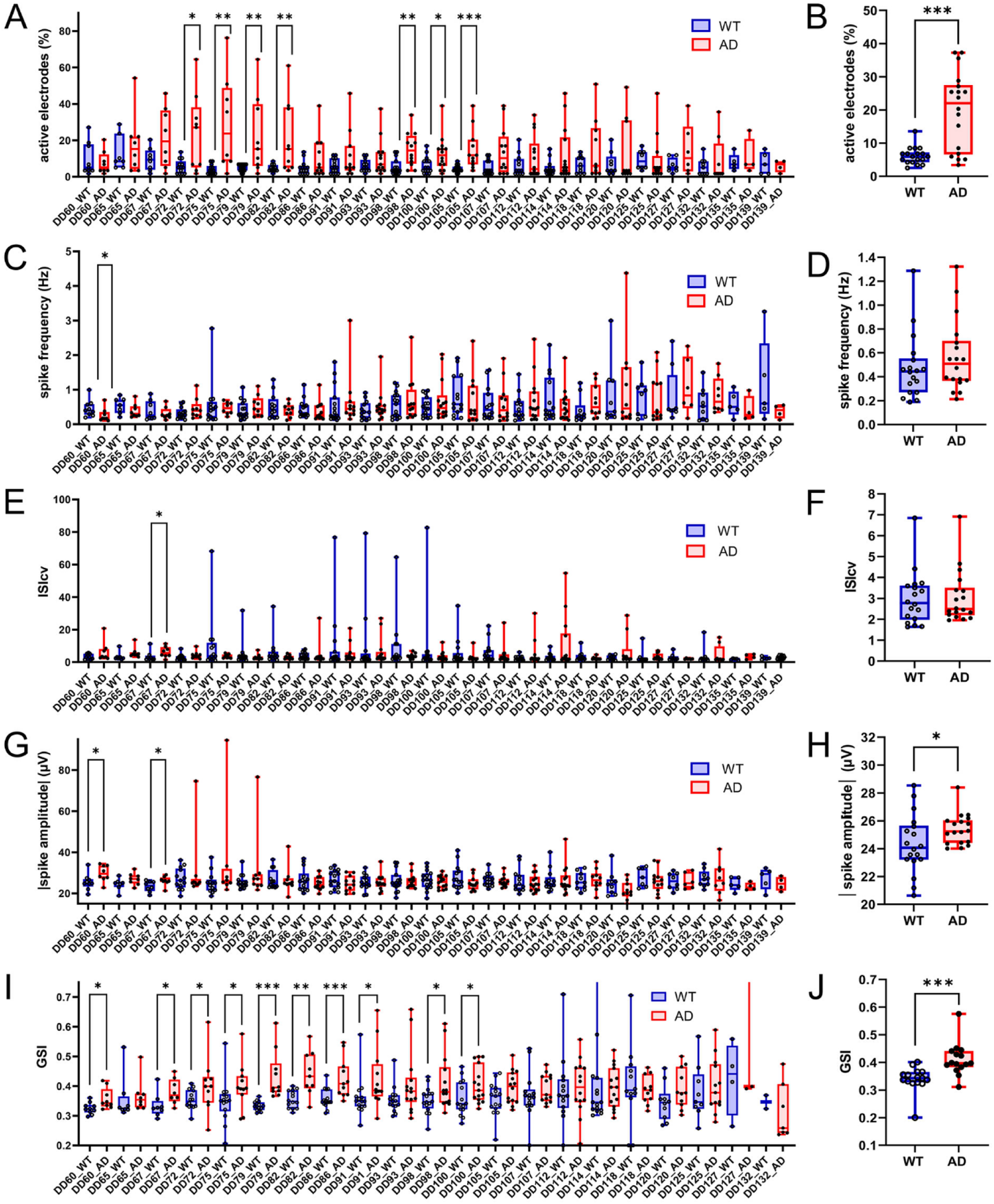
Differences in the spike activity in WT and AD cerebral organoids. **A, C, E, G, I:** Changes in spike parameters during the whole recording period, namely in the percentage of active electrodes (A), spike frequency (C), interspike interval coefficient of variation (ISIcv; E), spike amplitude (G), and global synchrony index (GSI; I). **B, D, F, H, J:** Average spike parameters during the 3 weeks of the highest electrical activity (*n*_WT_ = 16/3, *n*_AD_ = 15/3), namely the percentage of active electrodes (B), spike frequency (D), ISIcv (F), spike amplitude (H), and GSI (J); \**P* <0.05, \*\**P* <0.01, \*\*\**P* <0.001 (Mann-Whitney test).

Subsequently, we analysed burst parameters, which capture coordinated, high-frequency firing patterns that single-spike counts cannot, providing a more accurate measure of network-level activity and functional connectivity in organoids. The burst analysis revealed that AD organoids developed significantly more robust bursting behaviour over time (**Figure 4**). Compared to WT, AD organoids exhibited a significantly higher percentage of active electrodes and intraburst spike frequency (\**P* < 0.05, \*\**P* < 0.01 for both, and \*\*\**P* < 0.001 for the percentage of active electrodes; **Figure 4A**). Average burst parameters at the peak of the activity, including the percentage of active electrodes, intraburst spike frequency, intraburst spike number, and burst duration, were significantly higher in AD compared to WT organoids (\*\*\**P* < 0.001 in the first three parameters and \**P* < 0.05 in the burst duration; **Figure 4B**). While burst frequency and total burst count showed a slight trend toward higher values in WT, interburst interval trended higher in AD, and the interburst interval coefficient of variation (IBIcv) was similar across groups (**Figure 4B**). However, none of these differences were statistically significant. These observations imply that, while more neurons engage in bursts in AD organoids, the timing and frequency of bursts across the network remain essentially unchanged. Overall, burst parameters suggest that AD organoids exhibit enhanced temporal coordination of activity, along with increased recruitment of neurons to fire, consistent with greater network synchronisation.

**Figure 4:**
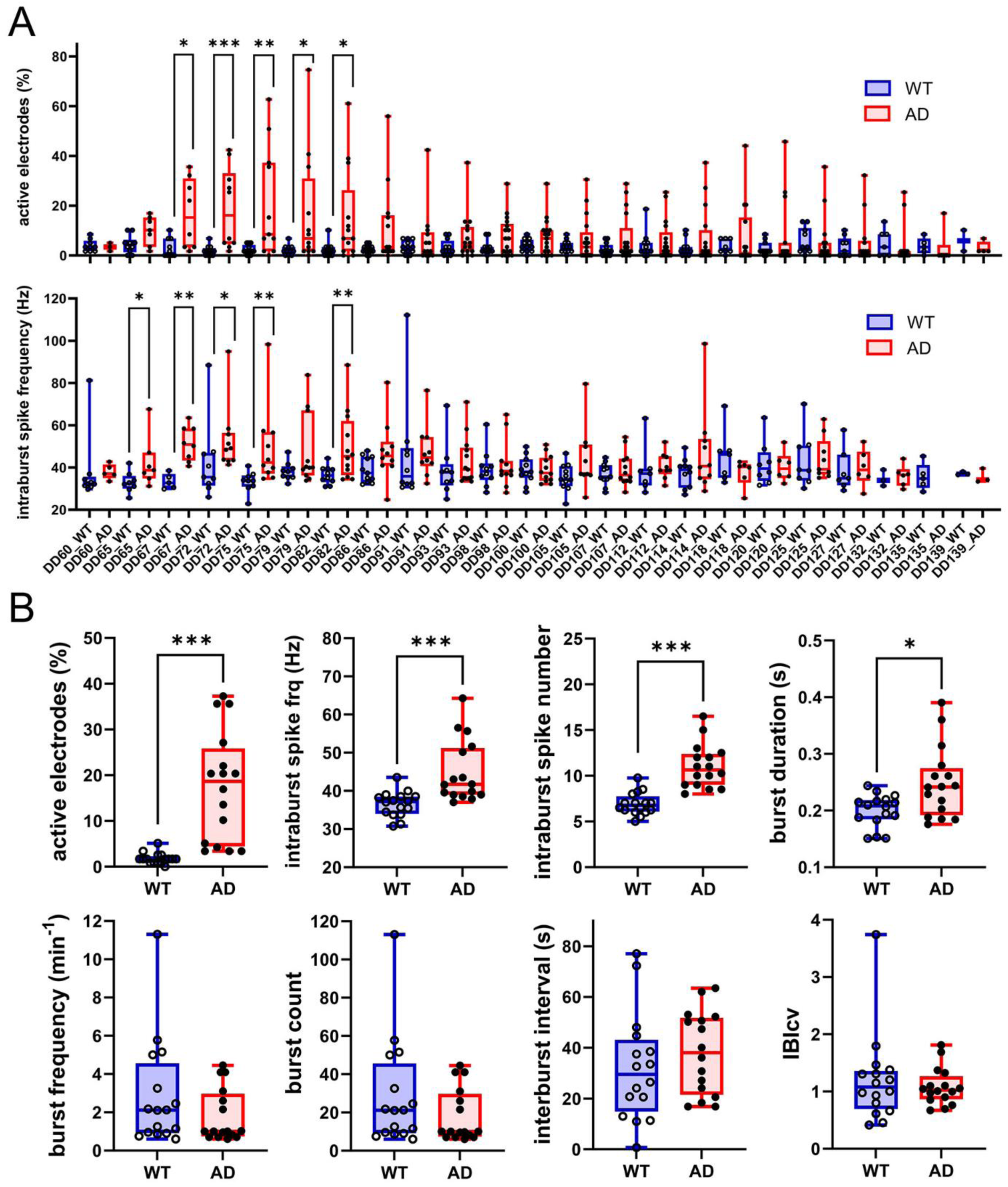
Differences in the burst activity in WT and AD cerebral organoids. **A:** Changes in the number of active electrodes (up) and intraburst spike frequency (down) during the whole approx. 2.5-months recording. **B:** Average burst parameters during the 3 weeks of the highest electrical activity (*n*_WT_ = 16/3, *n*_AD_ = 15/3) - a significant increase was observed in the number of active electrodes, intraburst spike frequency, intraburst spike number, and burst duration; IBIcv - interburst interval coefficient of variation; \**P* < 0.05,\*\**P* < 0.01, \*\*\**P* < 0.001 (Mann-Whitney test).

### 3.3 Aβ42/40 positively correlates with electrical hyperactivity in AD

To investigate whether Aβ pathology is linked to changes in neural network activity, we examined correlations between normalised Aβ42/40, normalised Aβ aggregate size, and normalised parameters of the electrical activity relative to the first time point in both WT and AD organoids. The normalised Aβ42/40, as well as the normalised Aβ aggregate size, was estimated in all included biological replicates at DD60, DD90, and DD130 (**Figure 1F**). These data were then used to examine possible coupling between normalised Aβ content and normalised electrical activity measures of the organoids, respecting the analogous time points. In this context, AD organoids demonstrated a significant positive correlation between Aβ42/40 and the percentage of active electrodes in spike activity and GSI (*r*_s_ = 0.94, \**P* < 0.05; and *r*_s_ = 1.00, \*\**P* < 0.01; **Figures 5A and 5B**). Also, a non-significant positive trend was observed in the case of the percentage of active electrodes in burst activity and intraburst spike frequency (*r*_s_ = 0.81, *P* = 0.09 for both; **Figures 5C and 5D**). Likewise, Aβ aggregate size displayed a non-significant positive trend with the percentage of active electrodes in spike activity, a significant positive correlation with GSI, non-significant correlation with the percentage of active electrodes in burst activity, and a significant positive correlation with intraburst spike frequency in AD organoids (*r*_s_ = 0.81, *P* = 0.09; *r*_s_ = 0.94, \**P* < 0.05; *r*_s_ = 0.74, *P* = 0.13; and *r*_s_ = 0.94, \**P* < 0.05; **Figures 5E, 5F, 5G, and 5H**). On the other hand, WT organoids showed no such correlation in both normalised Aβ42/40 and normalised Aβ aggregate size. Similar trends were apparent when we used the non-normalised data of the 3 individual biological replicates (**Supplementary Figure S1**). These findings suggest that, in AD organoids, elevated Aβ42/40 and Aβ aggregate size are potentially pathologically associated with increases in neuronal recruitment and network synchronisation, whilst this relationship is absent in WT organoids.

**Figure 5:**
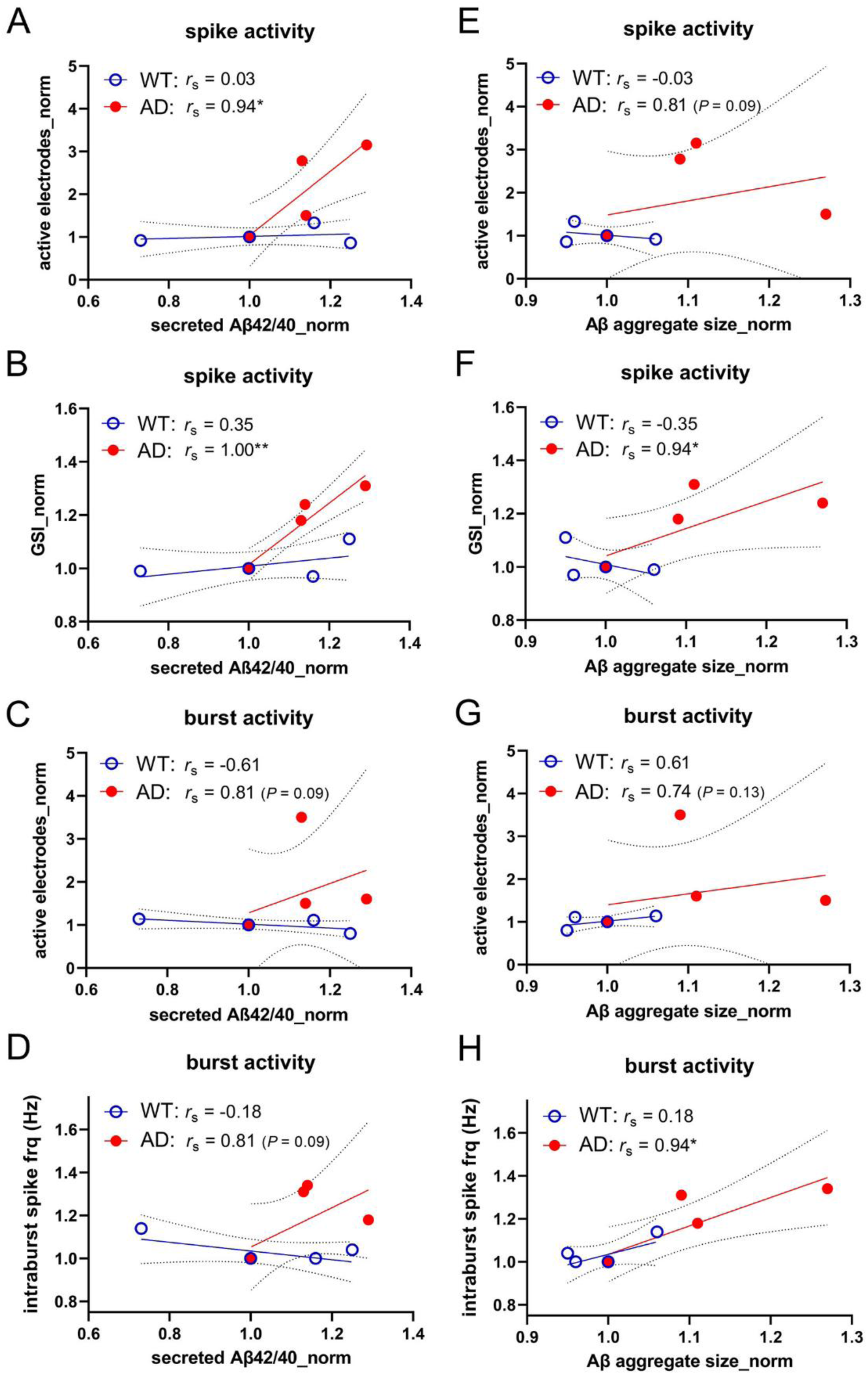
Correlation between the electrical activity and Aβ in WT and AD cerebral organoids. **A, B, C, D:** Correlation between Aβ42/40 and spike active electrodes (A, \**P* < 0.05), global synchrony index (GSI; B, \*\**P* < 0.01), burst active electrodes (C, *P* = 0.09), and intraburst spike frequency (D, *P* = 0.09). **E F, G, H:** Correlation between Aβ aggregate size and spike active electrodes (E, *P* = 0.09), GSI (F, \**P* < 0.05), burst active electrodes (G, *P* = 0.13), or intraburst spike frequency (H, \**P* < 0.05); *r*_s_ - the Spearman correlation coefficient. Data were normalised to the first timepoint, *i.e.* DD90/DD60 in the second and third differentiation, whereas DD130/DD90 in the first differentiation.

## 4. Discussion

In this study, we monitored spontaneous electrical activity in hiPSC-derived cerebral organoids carrying the fAD-causing *PSEN1* A246E variant over an extended maturation period (DD60–DD139). We found that AD organoids showed significantly elevated and more synchronised electrical activity compared to WT organoids, along with increased Aβ42 secretion and Aβ aggregation. Importantly, Aβ42/40 and Aβ aggregate size positively correlated with several measures of hyperexcitability and synchrony, indicating that Aβ dysregulation likely contributes to hyperexcitability and hypersynchrony in the early stages of AD.

### 4.1 Dynamic changes of electrical activity and hypersynchronous oscillations as specific features of early AD

Cerebral organoids generated from patient-specific hiPSC have proven valuable for modelling of fAD (Gonzalez *et al*., 2018; Arber *et al*., 2021; Yanakiev *et al*., 2022; Vanova *et al*., 2023). Unlike animal models, hiPSC-derived neuronal cultures preserve the human genetic background and natural developmental trajectories of neuronal networks, providing a relevant context for studying genotype–phenotype interactions (Shimada *et al*., 2022; Sreenivasamurthy *et al*., 2022). Recent studies have highlighted their utility for investigating AD-related processes such as Aβ deposition, synaptic dysfunction, and altered electrical activity (Ghatak *et al*., 2019; Lomoio *et al*., 2023). Particularly, several groups have reported early network hyperexcitability as a robust functional phenotype in AD organoids. For example, Yin & VanDongen (2021) revealed that cerebral organoids harbouring the *PSEN2* N141I variant displayed markedly enhanced neuronal excitability relative to isogenic control organoids. Similarly, Ghatak *et al*. (2021) demonstrated hypersynchronous discharges in hiPSC-derived organoids carrying *PSEN1* ΔE9, *PSEN1* M146V, and *APP*^swe^ variants, suggesting that such models could serve as functional platforms for testing anti-epileptiform interventions. Furthermore, Chen *et al*. (2021) showed that even sporadic AD (sAD) organoids exposed to serum, mimicking a leaky blood-brain barrier, exhibit disrupted excitatory dynamics in MEA recordings. Overall, these findings converge on the principle that dysregulated neuronal excitability and synchrony represent an early and conserved physiological hallmark of both fAD and sAD. This interpretation is additionally supported by computational modelling, which predicts network hyperexcitability as an early feature of AD pathophysiology (Stam & de Haan, 2024).

In this study, AD organoids displayed robust hyperexcitability and hypersynchrony, but only during the first half of the experiment; the activity progressively declined at later time points (**Figures 2, 3A, 3I, and 4A**). A similar temporal pattern has been documented across other organoid-MEA models. Although Ghatak *et al*. (2019) reported pronounced hyperexcitability in early-stage AD organoids carrying the *PSEN1* M146V variant (approximately DD42), Hurley *et al*. (2023) observed a shift toward hypoexcitability in organoids with the *PSEN1* L435F variant at later maturation stages (around DD180). Together with our findings in *PSEN1* A246E organoids, these studies suggest that the transition from hyper- to hypoexcitability may be a common feature across fAD variants as synaptic dysfunction and pathology accumulate. Complementary, in patients, seizures are more frequent during presymptomatic and prodromal stages of AD, whereas later phases are dominated by synaptic loss and cognitive decline (Targa Dias Anastacio *et al*., 2022; Lam *et al*., 2024; Sanchez-Rodriguez *et al*., 2024). Clinical EEG recordings reveal these early-stage electrical changes reflecting network hyperexcitability (Kamondi *et al*., 2024; Nous *et al*., 2024), while functional MRI and PET imaging indicate that early hyperconnectivity in specific cortical networks precedes major disconnection (Sperling, 2011; Abuwarda *et al*., 2025). Correspondingly, *APP-* and *PSEN-*transgenic mouse models show early hippocampal and cortical hyperexcitability, including hypersynchronous oscillations and epileptiform discharges, which appear before overt neuronal loss (Ziyatdinova *et al*., 2011; Bezzina *et al*., 2015; Ray *et al.,* 2024). These early changes were revealed by methods such as *in vivo* imaging and electrophysiological recordings (Busche *et al*., 2012; Kazim *et al*., 2017). Hence, the dynamic increase of electrical activity followed by its decrease, resulting in hypoexcitability, that we observed in AD organoids, is consistent with both human and animal studies and seems to be a characteristic feature of excitability changes during AD development.

### 4.2 Aβ correlates with the neuronal network hyperexcitability and hypersynchrony in early AD

Our data further demonstrated that, compared with WT, AD organoids showed a pronounced increase in Aβ42/40, accompanied by the formation of larger Aβ aggregates (**Figures 1E and 1F**), as was documented in previous studies (Ghatak *et al*., 2019; Yin & VanDongen, 2021), including studies working with cerebral organoids carrying the same *PSEN1* A246E variant as we did (Gonzalez *et al*., 2018; Vanova *et al*., 2023). Hernández-Sapiéns *et al*. (2020) showed that hydrogel-based 3D neuronal cultures generated from hiPSCs with the *PSEN1* A246E variant accumulate Aβ oligomers and deposits at very early stages (DD14), confirming that this variant robustly drives amyloid pathology. Extending these findings to another tissue context, Lavekar *et al*. (2023) reported that retinal organoids carrying the *PSEN1* A246E variant exhibit elevated Aβ42/40 along with increased phosphorylated tau (p-tau) immunoreactivity (around DD150), highlighting convergent AD-related changes across neural tissues. In contrast, tau pathology in our system (DD60-DD130) was sparse and showed no clear relationship with electrophysiological changes (not illustrated). This agrees with the prior report of Shimada *et al*. (2022) that p-tau phenotypes often emerge later or require additional stressors to become consistent. Consistently, clinical studies show that plasma Aβ42/Aβ40 becomes abnormal first, roughly 7–14 years before symptom onset, whereas plasma p-tau217 and p-tau181 rise several years later (Milà-Alomà *et al*., 2025). Animal models similarly reflect this sequence, with Aβ accumulation appearing first and tau pathology developing only months or even over a year later, once sufficient amyloid burden has accumulated (Metaxas *et al*., 2019). Taken together, these data suggest that amyloid-driven processes predominate in early network dysfunction, while p-tau may exert a greater influence at later disease stages. To summarise, in line with the amyloid cascade hypothesis (Selkoe & Hardy, 2016) and in agreement with the study by Hernández-Sapiéns *et al*. (2020), our results suggest that Aβ dysregulation could be the primary driver of early functional disturbances.

Finally, given that early Aβ accumulation precedes other pathological hallmarks, a key question is whether Aβ itself drives neuronal hyperexcitability and hypersynchrony. Evidence from animal studies indicates a causal link: opto- and chemogenetic interventions can prevent Aβ accumulation and restore calcium homeostasis, thereby normalising network activity and synaptic function (Yuan & Grutzendler, 2016; Kastanenka *et al*., 2017). Consistently, hiPSC-derived AD neurons and cerebral organoids exhibit synchronised firing and increased bursting associated with elevated Aβ dimers or oligomers (Ghatak *et al.,* 2019). In further support of this mechanism, Fernandez-Perez *et al*. (2021) demonstrated that intracellular Aβ oligomers enhance neuronal hyperexcitability *via* PKC-dependent synaptic mechanisms, promoting action potential spread across rodent primary hippocampal neurons without altering intrinsic membrane properties. Functional evidence also comes from targeted modulation of Aβ production. Inhibition of BACE1, the protease initiating Aβ generation from APP, reduces synapse numbers and suppresses synaptic transmission even in WT hiPSC-derived neurons, highlighting the sensitivity of human synapses to perturbations in APP–Aβ processing (Zhou *et al*., 2022). Lastly, Ng *et al*. (2022) revealed that hiPSC-derived cortical neurons from patients with early symptomatic AD exhibit Aβ-driven synapse dysfunction, which correlates with the clinical vulnerability of the same patients to Aβ burden. Together with our data showing positive correlations between Aβ42/40, Aβ aggregate size, and several measures of hyperexcitability and synchrony (**Figure 5 and Supplementary Figure S1**), these findings suggest that aberrant network activity in human AD-relevant systems may arise from early Aβ dysregulation, before the onset of neurodegeneration.

### 4.3 Conclusions

This study revealed a direct correlation between changes in Aβ content and electrical activity in AD cerebral organoids. The data demonstrate that the *PSEN1* A246E variant causes an imbalance in Aβ42/40 levels, which may be associated with transient hyperexcitability and hypersynchrony in AD cerebral organoids, supporting Aβ as an early disruptor of neural networks. The observed temporal dynamics of spontaneous electrical activity in AD organoids align well with the disease trajectory observed in patients and animal models, providing an explanation for the conflicting reports of hyper-versus hypoexcitability across studies. Our findings establish the therapeutic potential of focusing on Aβ-associated hyperexcitability in the early stages of AD and demonstrate the suitability of organoids as a translational model for mechanistic research and pharmacological testing.

## 7. Data Availability

The data will be made available upon a reasonable request.

## Author Contributions

Material preparation, data collection, and analysis were performed by Andrijana Angelovski, Hana Hribkova, Jiri Sedmik, Barbora Liscakova, Olga Svecova, Sona Cesnarikova, Katerina Amruz Cerna, Veronika Pospisilova, Martin Kral, Martina Kolajova, Petr Klimes, and Marketa Bebarova. The first draft of the manuscript was written by Andrijana Angelovski, Hana Hribkova, Jiri Sedmik, Barbora Liscakova, Dasa Bohaciakova, and Marketa Bebarova. All authors commented on previous versions of the manuscript and read and approved the final manuscript.

