## Supplementary Table S1 for "Patient-Derived *PSEN1* Cerebral Organoids Revealed Parallel Development of Amyloid-β Accumulation and Network Dysfunction"

### Supplementary Table S1: Media, primers, and antibodies.

#### Cerebral Organoid Neural Induction Medium (NIM)

| ITEM | COMPANY | IDENTIFIER | STORAGE CONCENTRATION | FINAL CONCENTRATION |
|---|---|---|---|---|
| DMEM/F12 | Thermo Fisher Scientific | Cat# 21331046 | Basal medium | Basal medium |
| N-2 Supplement | Thermo Fisher Scientific | Cat# 17502048 | 100X | 1X |
| GlutaMAX™ | Thermo Fisher Scientific | Cat# 35050061 | 100X | 1X |
| MEM Non-Essential AA | Thermo Fisher Scientific | Cat# 11140035 | 100X | 1X |
| Heparin sodium salt | Sigma | Cat# H3149 | 1 mg/mL (in $H_2O$) | 1 µg/mL |
| 2-Mercaptoethanol | Sigma | Cat# M3148 | 10mM (in PBS) | 0.05mM |
| ZellShield® | Minerva Biolabs | Cat# 13-0050 | 100X | 1X |

#### Cerebral Organoid Differentiation Medium without vitamin A (CODM−A)

| ITEM | COMPANY | IDENTIFIER | STORAGE CONCENTRATION | FINAL CONCENTRATION |
|---|---|---|---|---|
| DMEM/F12 | Thermo Fisher Scientific | Cat# 21331046 | Basal medium | 1/2 of Basal medium |
| Neurobasal™ | Thermo Fisher Scientific | Cat# 21103049 | Basal medium | 1/2 of Basal medium |
| N-2 Supplement | Thermo Fisher Scientific | Cat# 17502048 | 100X | 0.5X |
| Insulin | Sigma | Cat# I9278 | 10 mg/mL | 2.5 µg/mL |
| GlutaMAX™ | Thermo Fisher Scientific | Cat# 35050061 | 100X | 1X |
| MEM Non-Essential AA | Thermo Fisher Scientific | Cat# 11140035 | 100X | 0.5X |
| B-27™ Supplement minus vitamin A | Thermo Fisher Scientific | Cat# 12587010 | 50X | 0.5X |
| 2-Mercaptoethanol | Sigma | Cat# M3148 | 10mM (in PBS) | 0.1mM |
| ZellShield® | Minerva Biolabs | Cat# 13-0050 | 100X | 1X |

#### Cerebral Organoid Differentiation Medium with vitamin A (CODM+A)

| ITEM | COMPANY | IDENTIFIER | STORAGE CONCENTRATION | FINAL CONCENTRATION |
|---|---|---|---|---|
| DMEM/F12 | Thermo Fisher Scientific | Cat# 21331046 | Basal medium | 1/2 of Basal medium |
| Neurobasal™ | Thermo Fisher Scientific | Cat# 21103049 | Basal medium | 1/2 of Basal medium |
| N-2 Supplement | Thermo Fisher Scientific | Cat# 17502048 | 100X | 0.5X |
| Insulin | Sigma | Cat# I9278 | 10 mg/mL | 2.5 µg/mL |
| GlutaMAX™ | Thermo Fisher Scientific | Cat# 35050061 | 100X | 1X |
| MEM Non-Essential AA | Thermo Fisher Scientific | Cat# 11140035 | 100X | 0.5X |
| B-27™ Supplement | Thermo Fisher Scientific | Cat# 17504044 | 50X | 0.5X |
| 2-Mercaptoethanol | Sigma | Cat# M3148 | 10mM (in PBS) | 0.1mM |
| ZellShield® | Minerva Biolabs | Cat# 13-0050 | 100X | 1X |

#### Cerebral Organoid MEA cultivation medium

| ITEM | COMPANY | IDENTIFIER | STORAGE CONCENTRATION | FINAL CONCENTRATION |
|---|---|---|---|---|
| BrainPhys™ Neuronal Medium | STEMCELL Technologies | Cat# 05790 | Basal medium | Basal medium |
| N-2 Supplement | Thermo Fisher Scientific | Cat# 17502048 | 100X | 1X |
| B-27™ Supplement | Thermo Fisher Scientific | Cat# 17504044 | 50X | 1X |
| ZellShield® | Minerva Biolabs | Cat# 13-0050 | 100X | 0.5X |

#### qPCR primers

| TARGET | NCBI GENE ID | FW SEQUENCE 5’-3’ | RV SEQUENCE 5’-3’ |
|---|---|---|---|
| *GAPDH* | 2597 | AGCCACATCGCTCAGACAC | GCCCAATACGACCAAATCC |
| *GFAP* | 2670 | CCGACAGCAGGTCCATGT | GTTGCTGGACGCCATTG |
| *MAP2* | 4133 | TTGGTGCCGAGTGAGAAGA | GTCTGGCAGTGGTTGGTTAA |
| *NEFM* | 4741 | GAAATCGCTGCGTACAGAAAAC | TAATGGCTGTCAGGGCCTCTT |
| *RBFOX3* | 146713 | TACGCAGCCTACAGATACGCTC | TGGTTCCAATGCTGTAGGTCGC |
| *SOX2* | 6657 | TACAGCATGTCCTACTCGCAG | GAGGAAGAGGTAACCACAGGG |
| *TUBB3* | 10381 | TCAGCGTCTACTACAACGAGGC | GCCTGAAGAGATGTCCAAAGGC |

##### Antibodies

| ANTIBODY | COMPANY | IDENTIFIER | IHC DILUTION |
|---|---|---|---|
| β-Amyloid (D54D2) | Cell Signaling | Cat# 8243<br>RRID: AB_2797642 | 1:200 |
| CTIP2 | Cell Signaling | Cat# 12120<br>RRID: AB_2797823 | 1:200 |
| NeuN | Millipore | Cat# MAB377<br>RRID: AB_2298772 | 1:200 |
| MAP2 | Millipore | Cat# AB5543<br>RRID: AB_571049 | 1:200 |
| PAX6 | Cell Signaling | Cat# 60433<br>RRID: AB_2797599 | 1:200 |
| SOX2 | Cell Signaling | Cat# 3579<br>RRID: AB_2195767 | 1:200 |
| TUJ | Aves Labs | Cat# TUJ-0020<br>RRID: AB_2313564 | 1:200 |
| Hoechst 33342 Nucleic Acid Stain | Thermo Fisher Scientific | Cat# H3570<br>RRID: AB_3675235 | 1:1000 |
| Donkey Anti-Chicken AF647 | Jackson ImmunoResearch | Cat# 703-606-155<br>RRID: AB_2340380 | 1:400 |
| Donkey Anti-Mouse AF488 | Thermo Fisher Scientific | Cat# A-21202<br>RRID: AB_141607 | 1:400 |
| Donkey Anti-Rabbit AF568 | Thermo Fisher Scientific | Cat# A-10042<br>RRID: AB_2534017 | 1:400 |
| Donkey Anti-Rabbit AF647 | Thermo Fisher Scientific | Cat# A-31573<br>RRID: AB_2536183 | 1:400 |
