## Supplementary Figure S1 for "Patient-Derived *PSEN1* Cerebral Organoids Revealed Parallel Development of Amyloid-β Accumulation and Network Dysfunction"

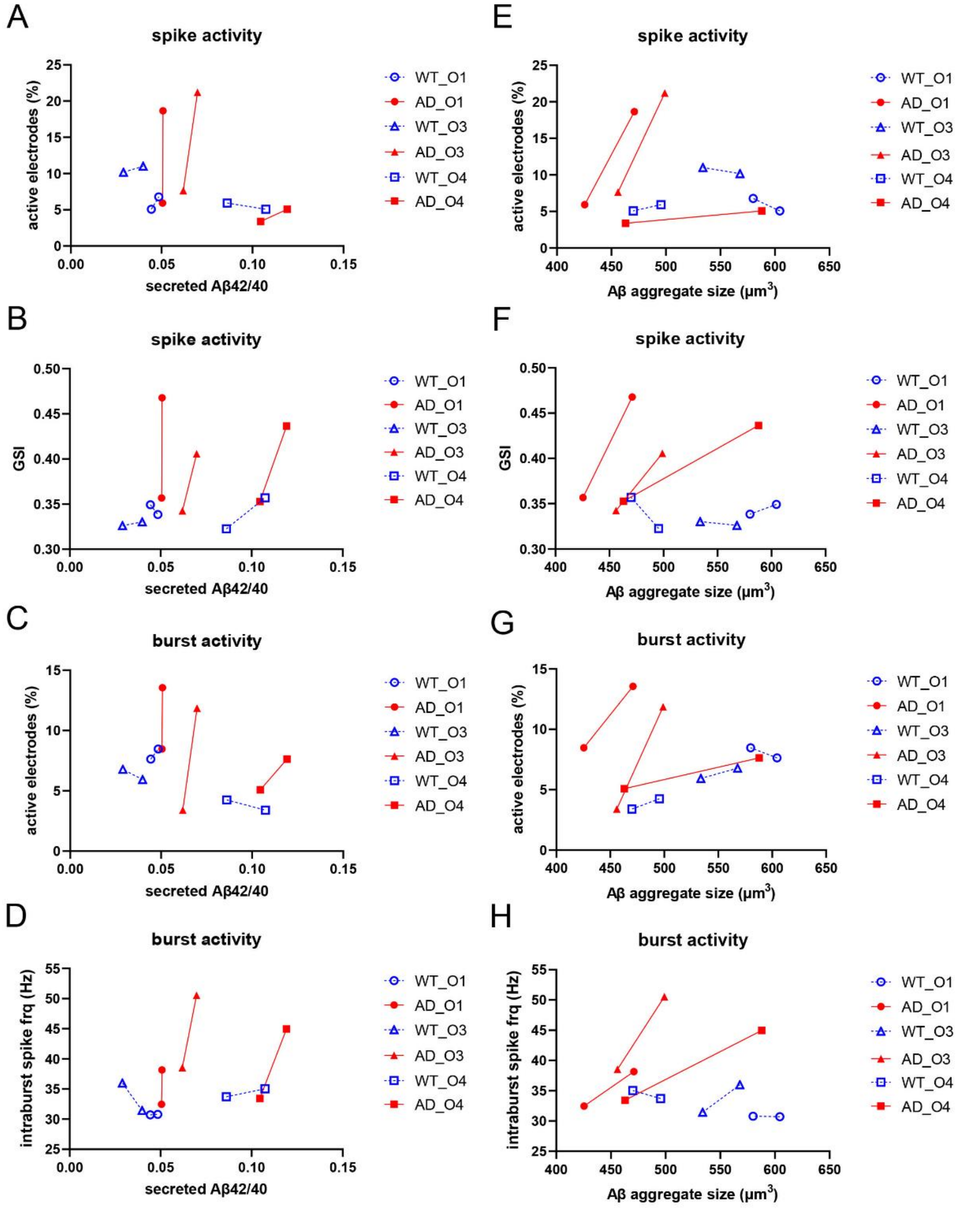


**Supplementary Figure S1: Correlation between the electrical activity and Aβ in WT and AD cerebral organoids from three independent differentiations - data in absolute values.**

**A, B, C, D:** Aβ42/40. **E, F, G, H:** Aβ aggregate size. Trends to a positive correlation can be mostly observed in AD (full red symbols), but not in WT (empty blue symbols), in both Aβ42/40 (A, B, C, D) and Aβ aggregate size (E, F, G, H).
